# Lorenz System in Thermodynamical Modelling of Leukemia Malignancy

**DOI:** 10.1101/034132

**Authors:** Igor Alexeev

## Abstract

Entropy rising within normal hematopoiesis is the core idea of the proposed thermodynamical model of malignancy in leukemia. Mathematically its description is supposed to be similar to the Lorenz system of ordinary differential equations for simplified processes of heat flow in fluids. The model provides description of remission and relapse in leukemia as two hierarchical and qualitatively different states of normal hematopoiesis with their own phase spaces. Phase space transition is possible through pitchfork bifurcation, which is considered as the common symmetrical scenario for relapse, induced remission and spontaneous remission of leukemia. Cytopenia is regarded as an adaptive reaction of hematopoiesis to entropy increase caused by leukemia clone. The following hypotheses are formulated: a) Percentage of leukemia cells in marrow as criterion of remission or relapse is not necessarily constant but a variable value; b) Probability of getting remission depends upon normal hematopoiesis reaching bifurcation; c) Duration of remission depends upon eradication of leukemia cells in induction or consolidation therapies; d) Excessively high doses of chemotherapy in consolidation might induce relapse.

## Foreword

Traditionally leukemia is considered as a unidirectional process of growth and expansion of leukemia clone into bone marrow. Resulting failure of normal hematopoiesis and involvement of other organs are regarded as complications of the disease. The corresponding model of leukemia as a proliferating tumor is based on its numerical aspect - amount of leukemia cells over time. Another aspect of leukemia – malignant effect of leukemia cells on normal hematopoiesis is often overlooked in modelling, despite the fact that it greatly determines the course of the disease and survival prognosis for the patient.

This article presents a model of leukemia malignancy as a process of entropy rising within normal haematopoiesis, caused by leukemia clone. Why thermodynamics is needed in description of leukemia and what to expect from it? Thermodynamics explains why something happens but does not show how it happens. It does not provide description of molecular mechanisms behind relapse or remission of leukemia, however it explains why spontaneous processes in leukemia are possible. Thermodynamics provides more accurate description of nonlinear aspect of leukemia dynamics than computational models of proliferating leukemia clone.

Mathematical description of entropy rising within normal hematopoiesis in leukemia is probably similar to entropy rising in heating shallow layer of fluid, known as Lorenz system. The role of leukemia clone in this context appears to be only a passive “heat” source while induced remission, spontaneous remission and relapse are considered as common manifestations of the same process of phase space transition through bifurcation in normal hematopoiesis.

The biggest weakness of this model is presentation of remission and relapse as two *qualitatively* different states of hematopoiesis in leukemia. The lack of experimental evidence for molecular mechanisms specific for remission or relapse can leave this model unsubstantiated. In addition, the model cannot be applied easily to analysis of solid cancers since the modelling conditions are different.

Thermodynamics provides a powerful tool for modelling different aspects of leukemia and thus it seems possible to build several thermodynamical models as alternatives to traditional linear model of leukemic clone expansion. Non-equilibrium aspect of leukemia might be modelled as a two-component unbalanced system of normal hematopoiesis and leukemic clone making a biological oscillator with entropy flow between its parts. A hybrid model of leukemia as a process of entropy rising within normal hematopoiesis and an upper level model of leukemia as a biological oscillator might be a next step in our modelling efforts.

## 1. Introduction.

### 1.1 Leukemia as a disease.

Leukemia is a group of cancers involving blood cells of bone marrow. Leukemia is the 11^th^ most common cancer worldwide, with around 352,000 new cases diagnosed in 2012 *[13].* Malignant transformation usually occurs at the level of a pluripotent stem cell or committed progenitor cells with more limited capacity for differentiation. It is generally accepted, that abnormally high proliferation rate and longevity lead to expansion of leukemic clone in bone marrow and often into various other organs and sites, such as liver, spleen, lymph nodes, central nervous system, kidneys and gonads. Resulting disruption of hematopoiesis cause anemia, infection, easy bruising or bleeding. In a typical case of acute leukemia such symptoms have usually been present for only days to weeks before diagnosis. Approximately 10^12^ leukemia cell have been estimated to be present by that time *[47]* which indicates that the growing leukemic clone coexisted with normal hematopoiesis for months without any apparent signs of its presence.

### 1.2 New mutations in leukemic clone as a basis of leukemia progression.

Relatively recent experimental evidence suggests that acute myeloid leukemia may originate from multiple clones of malignant cells *[9].* For chronic lymphocytic leukemia certain genetic events are found in the majority of cells which are considered as 'clonal driver mutations', whereas others, present only in a fraction of the tumor, are deemed to be 'subclonal driver mutations' *[43].* Sequencing studies revealed about 140 genes that when altered by intragenic mutations can act as driver mutations during tumorigenesis *[46].* The presence of sub-clonal driver mutations was associated with reduced survival in chronic lymphocytic leukemia *[24]*, and it seems that the degree of subclonality might serve as a cancer marker *per se.* Higher diversity is related to a higher mutation rate or longer tumor evolution with more replications *[41].*

### 1.3 Possible mechanisms of normal hematopoiesis disruption in leukemia.

Interaction between the healthy and cancer cell lines is often described through a competition for physical space resulting in an increased cellular degradation. This is consistent with the observation of an increase of markers for cell death such as lactate dehydrogenase *[3, 11, 22, 42]*. Several mechanisms underlying this spatial competition have been proposed: overcrowding which results in extinction of cells *[17]*; competition for a limited surface niche expressing certain receptors *[4, 48];* and apoptosis if no contacts to these receptors can be established *[15].* Other possible mechanisms include induction of cytopenia by impeding hematopoietic stem cells proliferation and differentiation *[26]* and competition for energy and nutrients *[39]*. Although molecular mechanisms of disruption are not known, at the level of cell populations hematopoiesis disruption is consistent with competitive exclusion principle (also known under several other names including Volterra-Lotka Law), which postulates that populations competing for the same limiting resource in one homogeneous habitat cannot coexist long enough *[18, 21]*. However, it is still debatable whether competitive exclusion principle developed for ecosystems can be applied for processes at cellular level.

### 1.4 Clinical remission and relapse as two states of hematopoiesis in leukemia.

The first manifestation of leukemia means not only expansion of leukemia clone into marrow and other organs but also disruption of normal hematopoiesis leading to severe complications of the disease. The goal of induction therapy of leukemia is attainment of a complete remission, which usually requires a period of marrow aplasia, or a “morphologic leukemia-free state,” following induction chemotherapy *[47].* Complete remission is currently defined as restoration of normal hematopoiesis with a blast cell fraction of less than 5% by light microscopic examination of the bone marrow. This relatively liberal definition reflects the difficulty of identifying leukemic blasts in regenerating marrow by morphologic criteria alone. Thus, patients with nearly 5% leukemic blast cells in their marrow specimens can harbor as many as 10^10^ leukemia cells *[5, 37].* Recurrence of leukemia, called relapse, after therapy is a common problem of leukemia. The goal of post-remission or consolidation therapy is to prolong complete remission by delaying or preventing relapse and to maximize the chance of cure *[47].* In a typical acute leukemia with chemotherapy the leukemic process is staged strictly as relapse or remission while correlations between the kinetic parameters of the normal and leukemic populations are suggested to characterize the leukemic state *[10].*

### 1.5 Spontaneous remission of leukemia.

Remission of leukemia without any specific therapy, called spontaneous remission, is an extremely rare and exceptional, relatively well documented but poorly understood phenomenon. Spontaneous remission of acute myeloid leukemia is almost always transient event, with a mean duration in the literature of 7.7 months (range 1–36) *[14].* In a typical case of spontaneous remission the full restoration of normal hematopoiesis and disappearance of blast cells occur in patient with acute leukemia and concurrent infection *[2, 7, 12, 16, 23, 27, 31, 32, 33, 36, 45]*, blood transfusion *[2, 14, 25, 36, 38]* or cytokine injection *[19, 44].* The underlying molecular mechanisms of spontaneous remission are still unknown. A potential role of bacterial or fungal infections and blood transfusions was suggested in spontaneous remission occurrence by triggering antileukemic and immune responses *[35].* Activation of cytotoxic T lymphocytes and macrophages in conjunction with an increased cytotoxicity of Natural Killer cells as well as cytokines of the immune system such as tumor necrosis factor, interferon gamma, and interleukin-2, released during infection may play a role in the occurrence of spontaneous remission *[6, 20, 29, 30, 31].* However, no clear link between spontaneous remission and infection or immune response was reported in at least one case *[8].* In another report spontaneous remission was detected after termination of pregnancy *[27].*

## 2. Lorenz system

Dynamical biological or physical systems display a variety of linear and nonlinear behaviors that can be described by corresponding mathematical models. Despite the diverse nature of processes, resulting mathematical description is quite similar, so it seems possible to understand some aspects of leukemia dynamics with the help of other models. This possibility of common mathematical description will be used to highlight similarities between leukemia and heat distribution in fluid flows.

Since leukemia increases entropy of normal hematopoiesis, Lorenz system is suggested to be used for modelling as it reflects a similar process of entropy rising in a uniformly heated and cooled shallow layer of fluid. Edward Norton Lorenz modeled weather and motion of air as it is heated and found that while being severely truncated version of Navier-Stokes equations (which arise from applying Newton's second law to viscous fluid motion), the system still preserves many of its characteristics. Detailed analysis of the Lorenz equations is out of scope of this article, however it is readily available and is commonly used as a simplified model of nonlinear system. Some terms of the theory of dynamical systems are used here when necessary in context of leukemia modelling. An introduction to the theory of dynamical systems is available [40].

In brief, Lorenz model for convection is the system of three ordinary differential equations (1),(2),(3):

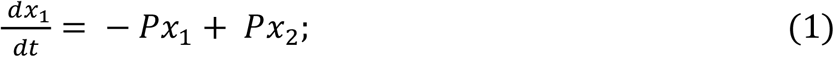

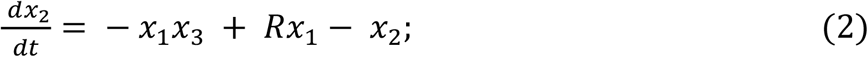

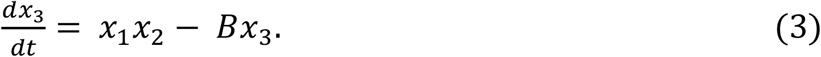

Given P=10 and B=8/3 (those are any of constant parameters) the system behavior depends upon R control parameter which reflects heating and relates to entropy of the system. For R<1 the origin is the only stable steady state (diagram 1). This situation corresponds to no convection in heating fluids. At R=1 there are two new stable steady states of the system where x(t)>0 and x(t)<0. These conditions correspond to the state of steady convection in fluids. There is also a pitchfork bifurcation, where a state transition between them is possible. The system remains stable until R=24.74 (diagrams 2 and 3).

**Diagrams 1-4.**
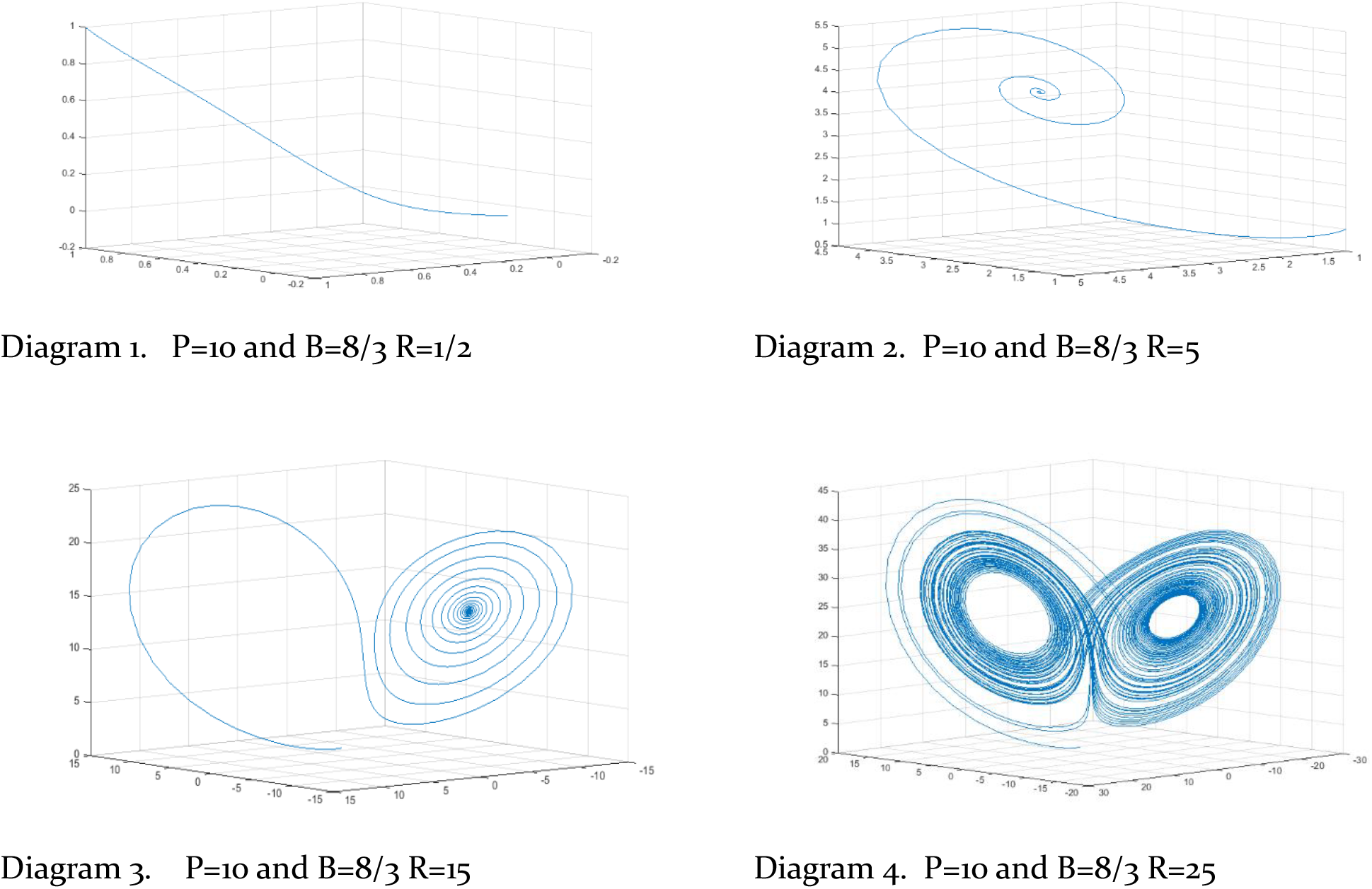
Lorenz attractor. From initial steady state (diagram 1) the system compartmentalizes (diagrams 2 and 3) making two new steady states and pitchfork bifurcation. With subsequent entropy increase the system loses stability (diagram 4).

Lorenz found that the system behaves chaotically for R>24.74 when it starts with a rotation around one of the focuses with an amplitude increasing with time, thereby forming a divergent spiral. After a number of such oscillations, the system suddenly goes toward the second available focus through a bifurcation and it continues an oscillatory motion around the second available focus along a divergent spiral. After a certain number of oscillations around this focus, the system jumps back to the vicinity of the previous focus, from which it again begins a new divergent oscillatory trajectory (diagram 4) (based on *[34]).*

## 3. Discussion

Leukemia is an incarnation of chaos. The chaos begins from one or several mutations in cell genome producing independently growing and quickly changing cell population of tumor. At certain moment it starts to affect the functioning of normal hematopoiesis probably competing with it for some common resources or indirectly disrupting its regulatory networks. Degree of disruption or malignancy determines survival prognosis for a patient with untreated disease. It is important to make distinction between the tumor *per se* and its disruptive effect on the normal hematopoiesis. If left untreated, a patient will be suffering because of failure of normal hematopoietic lineages caused by leukemia cells but not because of their presence.

Malignancy or harmful effect of leukemia clone on normal hematopoiesis can be expressed numerically through their approximate linear models. At the same time, in terms of thermodynamics harmful effect of leukemia on normal hematopoiesis can be described as an increase of its entropy, exceeding a certain physiological norm. These two approaches to description of leukemia demonstrate two aspects in its modelling: quantitative and dynamic. These aspects reflect different emphases and are not in conflict with one another. Quantitative or numerical aspect describes cellularity or cell composition of bone marrow as absolute and relative amount of cells. It reflects the stage of leukemia and the state of normal hematopoiesis. Dynamic aspect of leukemia modelling characterizes complex behavior of hematopoiesis in leukemia with transitions between relapse and remission as distinctive qualitative states of normal hematopoiesis. For our modelling purposes we use mathematical description of a similar process of entropy flow in a uniformly heated shallow layer of fluid known as Lorenz system. Similarities between two processes are not only entropy flows but also the same modeling conditions which are homogenous medium of fluid in Lorenz system and homogenous medium of bone marrow in leukemia.

Graphical solution of ordinary differential equations of Lorenz system is an abstraction which provides three clues for understanding leukemia. Firstly, at certain level of entropy the system compartmentalizes passing from an only stable state (diagram 1) to a new steady state with two available focuses within their phase spaces (diagrams 2 and 3). In our model of leukemia it indicates that upon appearance of leukemia clone normal hematopoiesis exists in two qualitatively different states - one state corresponds to remission and another to relapse. In other words, remission and relapse exist in leukemia as different states of hematopoiesis regardless of processes within tumor. Leukemia clone in this context is only a “heat” source which destroys normal hematopoiesis by increasing its entropy.

Idea of remission and relapse as two qualitatively different stable states of hematopoiesis in leukemia is derived directly from graphical solutions of Lorenz equations for R value greater than 15 (diagrams 3 and 4), however this assumption is still the biggest weakness of the proposed model, since no experimental evidence for any underlying molecular mechanisms in normal hematopoiesis is known. The best possible candidate for such a mechanism would be a signalling pathway for certain hematopoietic lineages which is active in remission and absent or clearly suppressed in relapse or *vice versa.*

Secondly, there is a pitchfork bifurcation between the phase spaces indicating an opportunity of swap between them. Bifurcation also indicates that remission and relapse are hierarchical states of hematopoiesis i.e. hematopoiesis can exist in one of the states but not in both of them simultaneously. Remission and relapse are the only two stable states of hematopoiesis in the model which can be observed clinically but pitchfork bifurcation is an unstable state of hematopoiesis. Finally, at a certain level of entropy the system loses stability making possible spontaneous phase space transition through the bifurcation (diagram 4). For our model of leukemia the chaotic behavior of Lorenz system corresponds to spontaneous remission in the course of the disease. Concurrent infection, blood transfusions or cytokine injections are factors contributing to stability loss and thus facilitating the spontaneous state transition. Spontaneous remission is usually transient since leukemia clone prevails.

Relapse of leukemia can be considered as another possible symmetrical scenario of spontaneous states swap. It involves reaching of bifurcation by hematopoiesis in the state of remission. Spontaneous remission and “spontaneous relapse” are the two symmetrical manifestations of the same process of the phase space transition through bifurcation. Successful course of chemotherapy should result in a period of marrow aplasia, which corresponds to bifurcation. Thus, the role of chemotherapy in leukemia appears to be dual since it is not limited only to eradication of leukemia clone. Chemotherapy has also a regulatory role driving normal hematopoiesis in relapse to bifurcation making possible state swap which results in remission.

According to the model, there are two ways of reaching remission in leukemia. Firstly, when normal hematopoiesis reaches bifurcation in relapse by chemotherapy or spontaneously. Such a remission can be of any duration depending on remaining leukemia clone or effectiveness of consolidation therapy. Secondly, an eradication of leukemia clone as a factor of entropy rise. In an ideal case of complete eradication it would mean complete cure of the disease with subsequent return of hematopoiesis to normal type. Relapse of leukemia can also be considered as a spontaneous event of reaching bifurcation but in the state of remission. While going further one can suggest that excessively high doses of chemotherapy in consolidation therapy could also induce relapse as state swap in remission.

Cytopenia observed at the first manifestation of leukemia can be interpreted as an adaptive reaction of hematopoiesis to entropy increase caused by leukemia clone [1]. It would be more appropriate to explain it by an idea of homeostasis as maintenance of thermodynamic temperature. With appearance of leukemia clone and subsequent increase of thermodynamic temperature of hematopoiesis, homeostasis of the whole system is maintained resulting in decrease of thermodynamic temperature of normal hematopoiesis. The latter leads to stem and progenitor cell proliferation and differentiation rate decrease and results in cytopenia. So, the normal hematopoiesis is suppressed in leukemia by a physiological mechanism which maintains homeostasis. With appearance of leukemia clone and at a certain level of entropy the normal hematopoiesis reaches bifurcation with subsequent phase space transition. Clinically this situation corresponds to the first manifestation of the disease or to relapse.

Thus, based on the model, some hypotheses can be formulated. Definition of remission and relapse solely through the state of normal hematopoiesis leads to the following: (a) percentage of leukemia cell in marrow for relapse or remission criterion is not necessarily constant but a variable value [1]; (b) probability of getting remission depends upon reaching bifurcation; (c) Duration of induced remission depends mainly upon degree of eradication of leukemia cells in therapy; (d) Excessively high doses of chemotherapy in consolidation therapy can induce relapse.

All four hypotheses can be tested in laboratory and by statistical analysis, however refinement of the model appears to be more difficult. Presumably, it should include integration of both computational and dynamic aspects of leukemia clone and normal hematopoiesis interaction by a model of leukemia as a biological oscillator in terms of non-equilibrium thermodynamics. The model verification would also require development of methods for entropy assessment in cell populations on laboratory models and in humans.

Regulatory role of thermodynamic potential on cellular processes was not largely discussed. It is known that at the cellular level high temperatures and some external shock factors upregulate expression of Heat Shock Proteins, - chaperons with mainly stabilizing and protective role within cells. Those changes might be also a part of regulatory effects of entropy fluctuations on normal hematopoiesis in leukemia, however, effects of weak thermodynamic temperature on cells and cell populations are still not proven.

Most existing models describe leukemia as linear process of tumor expansion in bone marrow. Resulting failure of normal haematopoietic lineages is regarded as complication of leukemia. Presented thermodynamical model considers leukemia as a non-linear process of entropy rising within normal haematopoiesis caused by leukemia clone. Mathematical description of these events is similar to Lorenz system of differential equations for heat flow in fluids and thus the following metaphoric conclusion seems appropriate: “Leukemia is fire in blood”.

## ASSOCIATED CONTENT

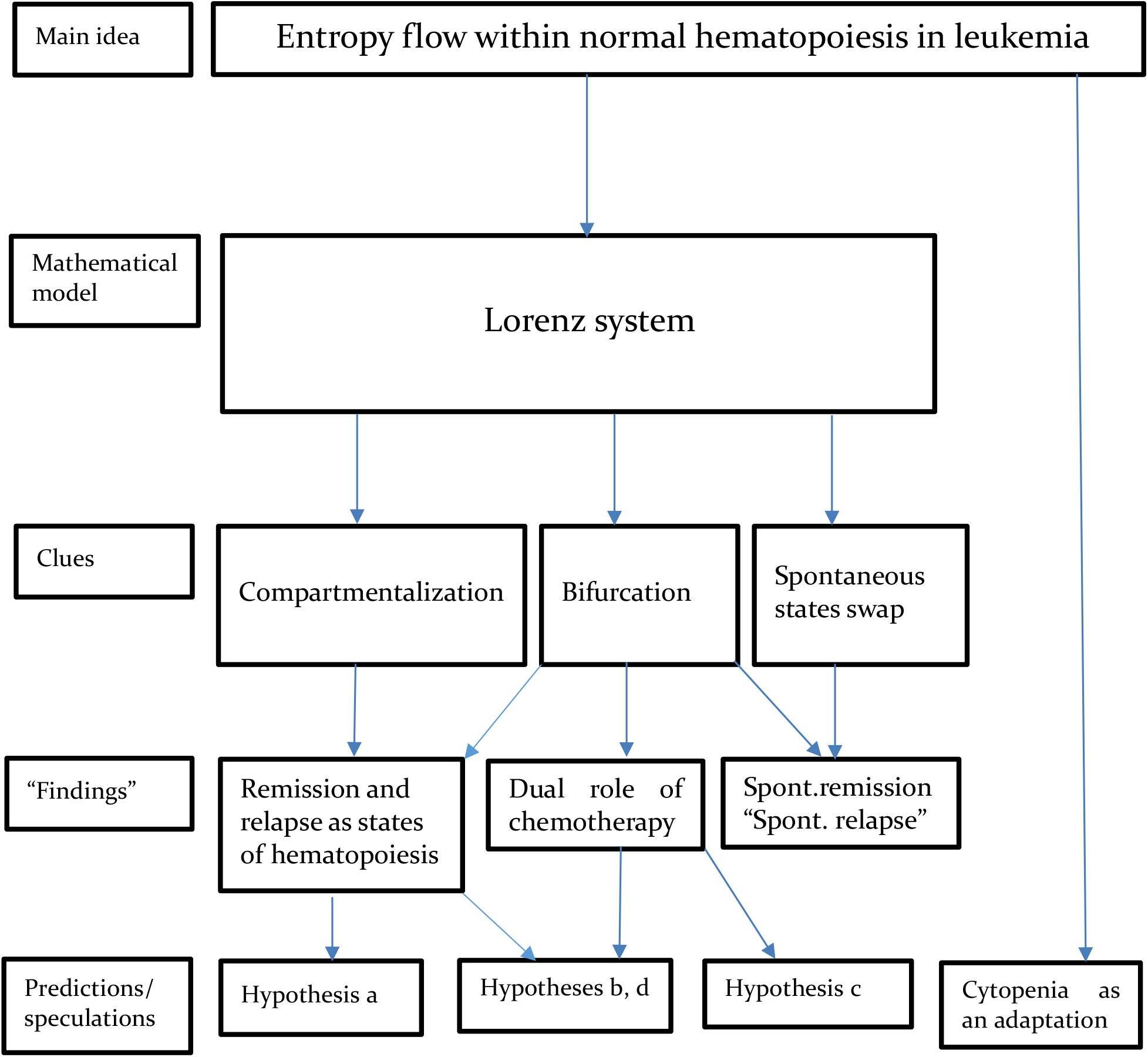
Structural diagram of major ideas in the article.

Not all connections are shown.

### The article in one sentence

“Remission and relapse are states of normal hematopoiesis but not percentage of blasts”.

MATLAB simulation scirpt for the Lorenz System equations in the time interval [0,100] with initial conditions [1,1,1] and r = 1/2.

~~~
clc
p=10;
b=8/3;
r=1/2;
f = @(t,a) [-p*a(1) + p*a(2); r*a(1) – a(2) – a(1)*a(3); -b*a(3) + a(1)*a(2)];
[t,a] = ode45(f,[0 100],[1 1 1]);
plot3(a(:,1),a(:,2),a(:,3))
~~~

